# A catalog of homoplasmic and heteroplasmic mitochondrial DNA variants in humans

**DOI:** 10.1101/798264

**Authors:** Alexandre Bolze, Fernando Mendez, Simon White, Francisco Tanudjaja, Magnus Isaksson, Ruomu Jiang, Andrew Dei Rossi, Elizabeth T. Cirulli, Misha Rashkin, William J. Metcalf, Joseph J. Grzymski, William Lee, James T. Lu, Nicole L. Washington

## Abstract

High quality population allele frequencies of DNA variants can be used to discover new biology, and study rare disorders. Here, we created a public catalog of mitochondrial DNA variants based on a population of 195,983 individuals. We focused on 3 criteria: (i) the population is not enriched for mitochondrial disorders, or other clinical phenotypes, (ii) all genomes are sequenced and analyzed in the same clinical laboratory, and (iii) both homoplasmic and heteroplasmic variants are reported. We found that 47% of the mitochondrial genome was invariant in this population, including large stretches in the 2 rRNA genes. This information could be used to annotate the mitochondrial genome in future studies. We also showed how to use this resource for the interpretation of pathogenic variants for rare mitochondrial disorders. For example, 42% of variants previously reported to be pathogenic for Leber Hereditary Optic Neuropathy (LHON) should be reclassified.

## Introduction

Mitochondrial diseases are among the most common of inherited disorders, with an estimated combined prevalence of 1 in 5,000 (Gorman et al., 2015; Schaefer et al., 2008). Mitochondrial disorders can be caused by variants encoded in nuclear (nDNA) or mitochondrial DNA (mtDNA); we focus here specifically on mtDNA variants. The mitochondrial genome codes for 13 protein-coding genes, 22 transfer RNA (tRNA) genes, 2 ribosomal RNA (rRNA) genes, and the non-genic displacement (D)-loop (Anderson et al., 1981). Unlike the nuclear genome, there are no introns, and there are very few non-coding bases in between genes. Human mitochondria are inherited through the maternal line, and there are multiple copies (>>2 copies) in every cell. Mitochondrial DNA can be uniform in sequence (homoplasmic) or can have variable sequences (heteroplasmy) within an individual cell. The level of heteroplasmy (or the proportion of mutated and wild-type mitochondria in a cell) can vary over time and between tissues (Wei et al., 2019). It is important to assess the level of heteroplasmy because often a phenotype is observed only if the levels of mutated mitochondrial DNA reach a certain level, deemed the “threshold effect” (Russell et al., 2020). Reports show that at the cellular level, a phenotype is typically observed if the heteroplasmy levels are above 70% (Russell et al., 2020), although some variants have been reported to have a phenotypic impact at levels as low as 20%. At the level of an individual, differences in levels of heteroplasmy can lead to varying phenotypic presentations of the same disease (Chinnery and Samuels, 1999; Chinnery et al., 2002).

The analysis of genetic variation in the population has been an efficient tool to understand the role and essentiality of genes and functional domains. It is also an essential tool to assess the pathogenicity of variants underlying rare disease. For example, scientists have recently drawn maps of constrained coding regions using the Genome Aggregation Database (gnomAD) (Havrilla et al., 2019), which highlighted regions depleted of non-synonymous variants across a large adult population. These regions pointed to genes and functional domains (including some without any known function) that may cause severe developmental phenotypes when mutated (Havrilla et al., 2019). Another example is the development of a framework that uses population allele frequency information to assess whether a variant is “too common” to be pathogenic for a specific disease, given the prevalence of this disease in the population and its assumed genetic architecture (Whiffin et al., 2017). These two examples illustrate how large databases such as Bravo (University of Michigan and NHLBI, 2018) and gnomAD (Karczewski et al., 2019) have been used to aid the interpretation of the human genome. However, these two databases do not have information on variants in the human mitochondrial genome (last checked in March 2020).

Before the public release of this study, MITOMAP (Lott et al., 2013) and HmtDB (Preste et al., 2019) were the two largest publicly available databases of human mtDNA variants, and have been used to assess the pathogenicity of mtDNA variants (Richards et al., 2015). For example, MITOMAP was used to select candidate mtDNA variants that may cause tubulointerstitial kidney disease (Connor et al., 2017). However, both databases gathered mtDNA variant information drawn from nearly the same ∼50,000 full mitochondrial genomes reported in GenBank, resulting in three known and reported limitations (Richards et al., 2015; Wong et al., 2020). Firstly, these databases are affected by biases in recruitment and are enriched for samples derived from patients with inherited mitochondrial disease. Secondly, mitochondrial genome sequences uploaded in GenBank come from different sources and are of unequal (and unknown) quality. These biases likely skew baseline rates of variation and estimates of allele frequencies. Lastly, these databases did not include heteroplasmic variants, which are essential when studying mitochondrial disorders (Wallace, 2018).

Here we provide a research resource of all mtDNA variants identified in 195,983 unrelated individuals without bias towards individuals with a mitochondrial disorder. All of the mitochondrial genomes were sequenced in the same clinical laboratory and analyzed using the same mitochondria-tailored pipeline. This resource includes both homoplasmic and heteroplasmic variants. After the characterization of mitochondrial DNA variation in humans, we report on two direct applications of this resource. We first studied the constraint on the mitochondrial genome sequence, pinpointing the most constrained regions. This will enable new annotations of particular relevance in the rRNA and tRNA genes. We then evaluated the utility of this resource for the interpretation of disease-causing variants by analyzing those reported to be pathogenic for Leber’s Hereditary Optic Neuropathy (LHON, OMIM:535000), the genetics of which has been studied for the past 30 years (Wallace et al., 1988).

## Results

### Creation of a high-quality catalogue of variation for the mitochondria

To create a resource that could be used to study rare mitochondrial disorders, we needed to aggregate information from a population who was not enriched in patients with mitochondrial disorders. Here, we sequenced individuals who are Helix users. There were no inclusion or exclusion criteria based on a mitochondrial disorder. The only inclusion criteria were: (i) being 18 years old or more, (ii) living in the United States at the time of consent, and (iii) having a unique email address. For all individuals, we sequenced their Exome+^®^, which includes the sequence of the full mitochondrial genome, followed by analysis of the mitochondrial genome using a mitochondria-specific pipeline (**Methods**). We then performed several quality control steps including the standard clinical laboratory analysis for the nuclear genomes: (i) quality of the overall sequencing output, (ii) assessment of contamination levels and re-collection of contaminated samples, and (iii) sex matching (**Figure 1A**). We also filtered samples that had five or more heteroplasmic variants outside of the hypervariable region as having more was considered very unlikely to be the result of true heteroplasmy, and more likely to be due to very low levels of contamination (originating from food in most of these cases) (**Figure S1**). Lastly, for all individuals, we calculated (i) ancestry using ADMIXTURE, (ii) mitochondria haplogroups using Haplogrep, and (iii) relatedness using Hail’s `pc_relate` function. We removed second-degree or closer related individuals. After applying these steps, we had 195,983 individuals and mitochondrial genomes to analyze and aggregate (**Figure 1A-C**). While almost all lineages present in the most recent version of PhyloTree (van Oven and Kayser, 2009) were represented in our dataset, 91.2% of the haplogroups were part of the Eurasian N lineages (**Figure 1D, Table S1**). The median age group was 46-50 years old (**Figure 1E**) and were 52.3% female.

**Figure 1.**
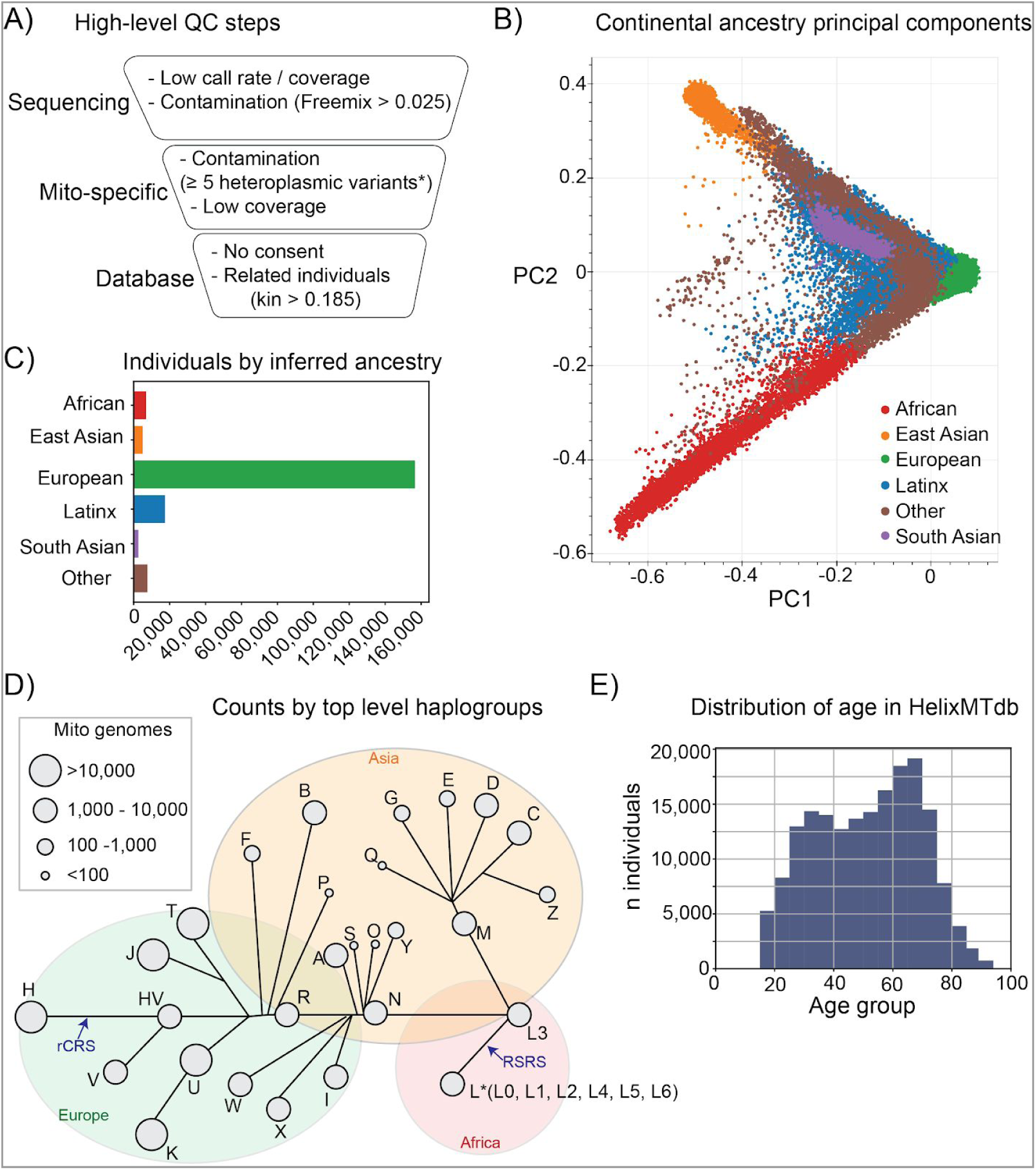
Overview of the 195,983 individuals and mitochondrial genomes aggregated in HelixMTdb. **(A)** Overview of the different quality control steps undertaken to narrow down the list of individuals included in this resource. **(B)** Continental ancestry principal components. Ancestries were inferred from coefficients from ADMIXTURE analysis. The majority of individuals in the “Other” category are individuals that are admixed. **(C)** Number of individuals per inferred continental ancestry. **(D)** Distribution of top-level haplogroups represented in HelixMTdb. Black lines define the mitochondrial phylogenetic tree originating at the mitochondrial genome RSRS. rCRS refers to the revised Cambridge sequence, which is the mitochondrial genome reference in this study. This figure was adapted from a figure on the Mitomap website under a Creative Commons Attribution 3.0 license. **(E)** Age-group distribution of individuals included in the resource. Age-group was self-reported at the time of providing a saliva sample. Individuals aged above the age 89 were all grouped together.

The next step was to assess the quality of the variant calls used to create the dataset. The mean base coverage was 182x. We first filtered out poor quality calls (GQ < 21 or DP <10). Next, we compared the allele frequencies of variants in HelixMTdb and MITOMAP as a quality check for the accuracy of our calls. We first assessed variants in a three small and known hard-to-sequence regions (Andrews et al., 1999; Wei et al., 2019): bases 300-316, 513-525, and 16182-16194. The majority of variants in these regions were insertions or deletions, and only a minority of these variants were observed in both datasets (**Figure S2**), which reinforced the fact that these regions are hard to sequence. We decided to filter out variants in these regions and not represent them in the final HelixMTdb. However, the absence of variants in these regions does not imply that these positions are not variable in the population. After removing the variants in known hard-to-sequence regions, the concordance of allele frequencies between the two databases was high (rho=0.82, Spearman Rho) (**Figure 2A, Figure S2**). The majority of variants unique to HelixMTdb (**Figure 2B**) and unique to MITOMAP are singletons in each, respectively (**Figure 2C**). These results provided confidence in the accuracy of the calls generated by our mitochondria-specific bioinformatics pipeline. On average, individuals had 25 homoplasmic variants (range: 0-99, median: 27), and 1 heteroplasmic variant (range: 0-13, median: 0) (**Figure 2D-E**). The number of homoplasmic variants per individual is explained mostly by the haplogroup and the evolutionary distance of that haplogroup from the haplogroup of the reference genome (rCRS). All individuals with haplogroups from the L and M lineages carried at least 22 homoplasmic variants (**Figure 2D**). There was also a small but significant increase of haplogroups in the L and M lineages among individuals with at least 1 heteroplasmic variant (L vs N: OR=1.7 p=4.2E-108, Fisher’s exact test; and M vs N: OR=1.3, p=1.5E-27, Fisher’s exact test) (**Figure 2E**). After aggregation of these results per variant over the sequenced population, we find a set of 14,324 mtDNA variants observed in at least one individual.

**Figure 2.**
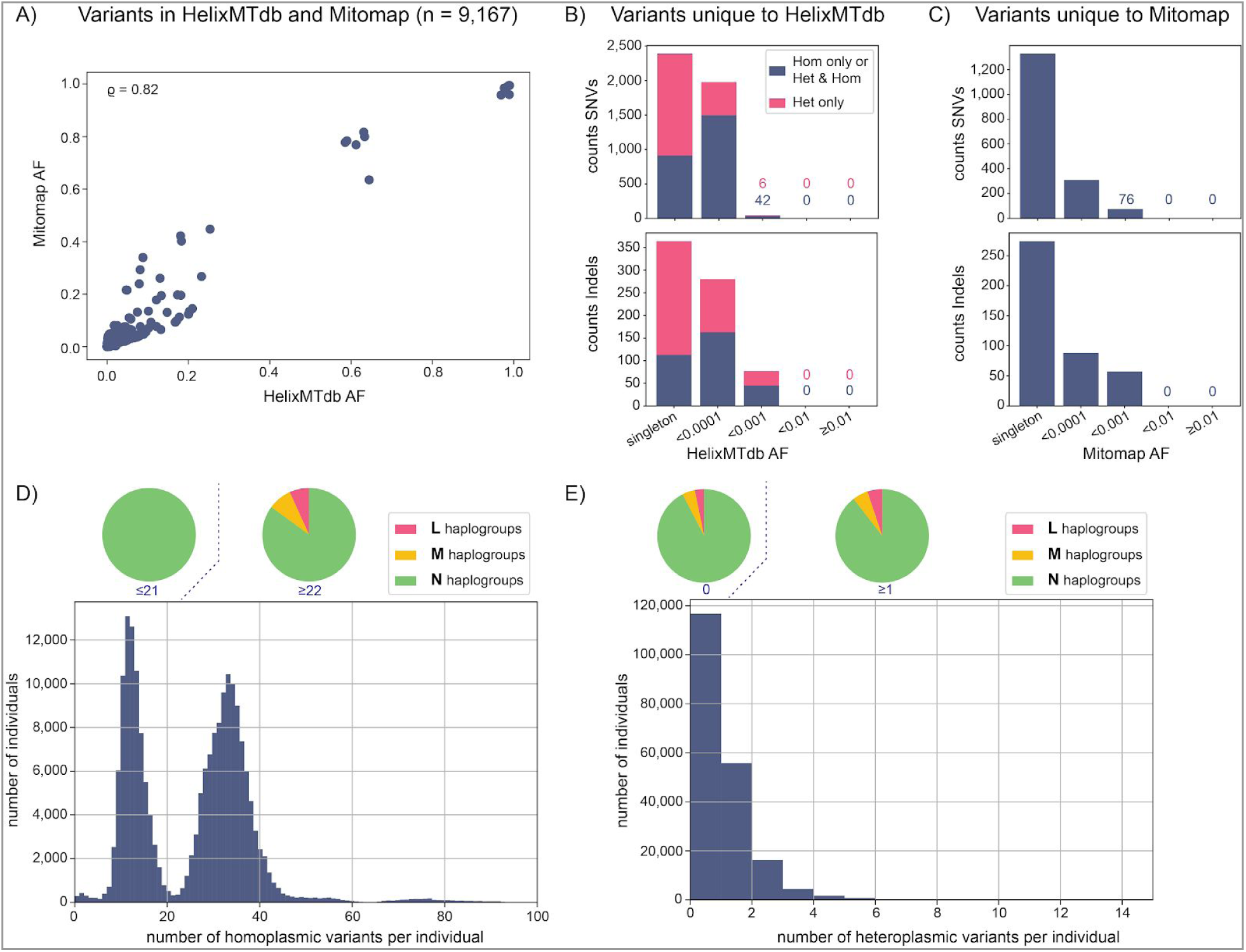
Reproducibility and accuracy of the variant calls. **(A)** Comparison of allele frequencies (AF) for variants present in both HelixMTdb and MITOMAP. P is the Spearman rho. **(B)** Allele frequencies (AF) of variants unique to HelixMTdb. The top panel represents SNVs and the bottom panel insertions and deletions. Of note, the majority of singletons unique to HelixMTdb are variants only observed at heteroplasmic levels in the population (represented by a pink box). **(C)** Allele frequencies (AF) of variants unique to MITOMAP. Top panel is for SNVs and the bottom panel for insertions and deletions. Mitomap does not report on heteroplasmic calls/variants. **(D)** Distribution of number of homoplasmic variants per individual. The pie charts represent the distribution of haplogroup lineages in individuals with a low number (less than the median) of homoplasmic variants, and in individuals with a high number (more than the median) of homoplasmic variants. **(E)** Distribution of number of heteroplasmic variants per individual. The pie chart represents the distribution of haplogroup lineages in individuals with a low number (n=0) of heteroplasmic variants, and in individuals with a high number (n≥1) of heteroplasmic variants. L, M and N haplogroups refer to all haplogroups downstream these 3 key nodes in the Phylotree (**Figure 1D** and **Table S1**).

The full database, HelixMTdb, can be downloaded (link in **Data availability** section, or https://Helix.com/Mito). It has information on the identity of variants, the number of times they were observed as homoplasmic or heteroplasmic, and the haplogroup(s) on which the variants were found.

### Characterization of mitochondrial DNA variation in 195,983 individuals

The distribution of allele frequencies of variants across the mitochondrial genome is represented in **Figure 3A**. The majority of variants (66%, n=9,400) were found to be present in less than 1 in 10,000 individuals in this cohort, with 24% of the variants (n=3,385) observed in only one individual (**Figure 3B**). Only 0.2% of the variants (n=35) were present in more than 10% of the individuals. We identified 13,435 single nucleotide variants (SNVs), 651 insertions, 237 deletions, and 1 indel (**Figure 3C**), and observed a higher abundance of transitions (73% of unique SNVs) than transversions (**Figure 3D**). Lastly, 51% of the variants (n=7,303) were observed both as homoplasmic and heteroplasmic in the population, whereas 29% of the variants (n=4,188) were only observed in homoplasmic calls, and 20% of the variants (n=2,833) were only observed in heteroplasmic calls (**Figure 3E**). For heteroplasmic variants, we defined the Alternate Read Fraction (ARF) to quantify the level of heteroplasmy observed. The distribution of the mean ARF of heteroplasmic calls for variants only seen at heteroplasmic levels is skewed towards lower ARF compared to the distribution for variants seen at homoplasmic and heteroplasmic levels in the population (**Figure 3F).**

**Figure 3.**
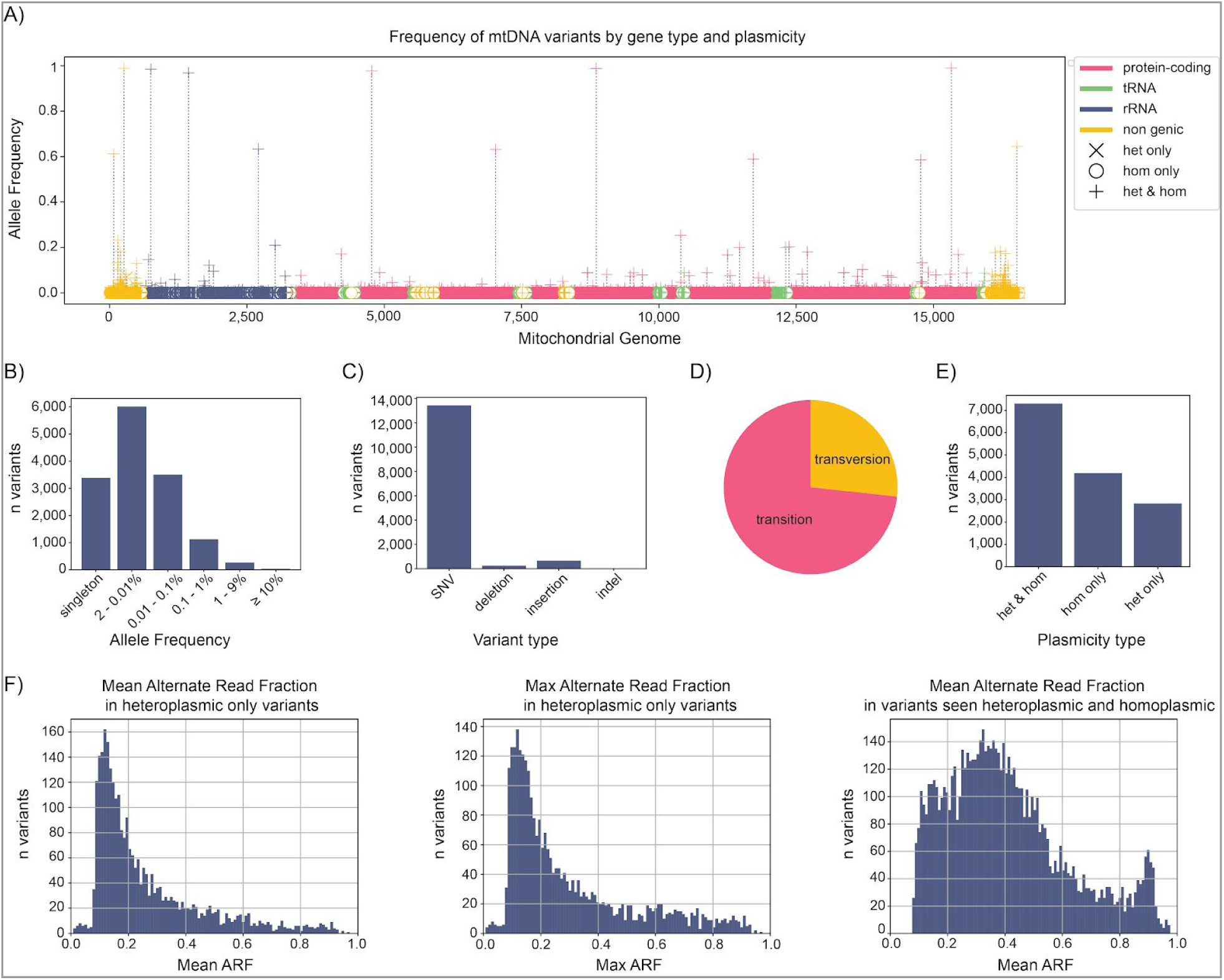
Characterization of the 14,324 variants identified in the population. **(A)** Linearized view of mitochondrial genome. pink: protein-coding genes; green: tRNA genes; blue: rRNA genes; yellow: noncoding. Lollipops above genomic features indicate variants observed at heteroplasmic levels only (x), at homoplasmic levels only (o), and at both heteroplasmic and homoplasmic levels (+) of plasmicity. **(B)** Variants were grouped by their frequency in this cohort. **(C)** Counts by variant type are indicated. **(D)** Proportion of transition, transversion. Analysis is restricted to SNVs and bi-allelic variants. **(E)** Distribution of variants that are seen in HelixMTdb only at homoplasmic levels (hom only), only at heteroplasmic levels (het only), or both (hom & het). Present in both means that there is at least one occurrence of the variant as homoplasmic and one occurrence as heteroplasmic for the given variant. **(F)** Distribution of the mean (left panel) and max (center panel) Alternate Read Fraction (ARF) for variants seen at heteroplasmic levels only. The right panel shows the mean ARF for variants seen at both homoplasmic and heteroplasmic levels in the population.

### The mitochondrial genome is not tolerant to truncating variants in protein-coding genes

We hypothesized that variants predicted to be damaging are less frequent than non-damaging variants in the general population, and are more often heteroplasmic (higher ratio (heteroplasmic calls) / (heteroplasmic + homoplasmic calls)). Of the 9,607 unique variants in protein-coding genes, only 85 (0.9%) were putative loss-of-function (LoF): 48 frameshift variants, 27 stop-gained variants, and 10 stop-loss variants (**Table S2)**. They were found in all genes (**Figure 4A**). These LoF variants were extremely rare in the population, with a mean allele frequency 0.007%, which is far lower than 0.14%, the mean allele frequency of all variants in protein-coding genes. Moreover, there was a significant enrichment of heteroplasmy among calls for predicted LoF variants: heteroplasmies represent 23% of all the calls for these variants, compared to ∼1% of the calls for variants predicted to be of medium or low severity (26% vs 1%; p=2.2E-240, Fisher’s exact test) (**Table S2**). In particular, all 48 frameshift and 26/27 stop-gained variants were only observed in the heteroplasmic state in the population at low levels of heteroplasmy (the average max ARF observed across these variants was 0.15) (**Figure 4B**), suggesting that they may not be tolerated when homoplasmic.

**Figure 4.**
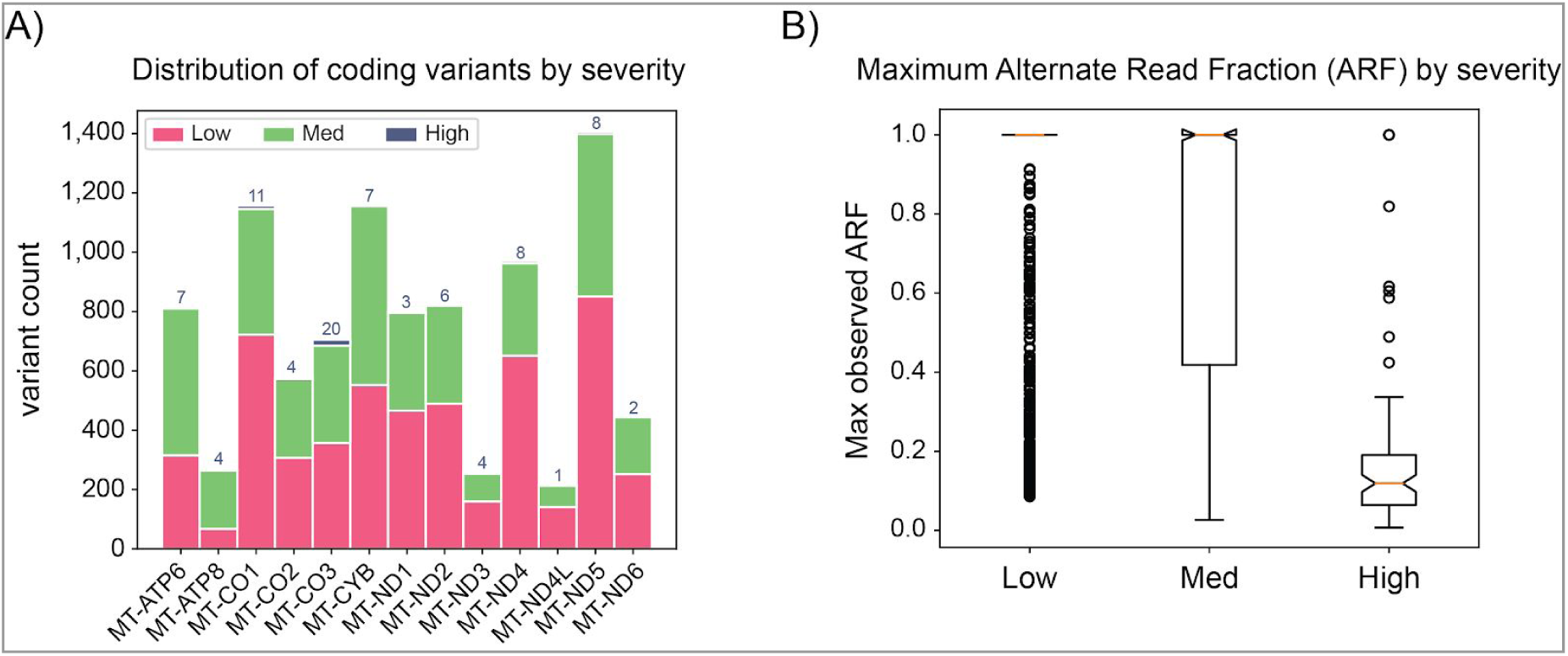
Intolerance to loss-of-function variants in protein-coding genes. **(A)** Summary counts of variants per gene, colored based on predicted severity. Pink: low; green: med; blue: high; yellow: unknown. Severity was annotated using VEP most_severe_consequence and grouped as follows: high (stop gained, frameshift, stop lost), medium (nonsynonymous, inframe indel, coding sequence variant, protein altering variant), low (synonymous, incomplete terminal codon variant). **(B)** Distribution of the maximum observed Alternate Read Fraction (ARF) in protein-coding genes for each severity category.

The only nonsense variant observed at homoplasmic levels was p.M1* in the *MT-ND1* gene. A few lines of evidence suggested that p.M1* in *MT-ND1* may not be a true loss-of-function: (i) it was observed in 89 individuals at homoplasmic levels, and none at heteroplasmic levels, (ii) it was only observed on haplogroup T, suggesting that this is a common polymorphism in a specific haplogroup, (iii) it was also observed in 6 individuals in MITOMAP, all of them belonging to the T1a haplogroup, (iv) there is a common missense variant at this position (m.3308T>C), and (v) the next methionine is at amino acid position 3, which may serve as an alternate start codon. These results indicate that all protein-coding genes in the mitochondrial genome were highly intolerant to LoF variants, especially at homoplasmic levels.

Outside of the protein-coding genes, we also observed intolerance to variants predicted to be damaging, especially in the 22 tRNA genes where variant annotations exist. There were 1,046 unique variants that mapped to the 22 tRNA genes. We classified the predicted pathogenicity of each tRNA variant using the scoring model from MitoTip (Sonney et al., 2017). There were 84 (8.0%) observed variants classified as known (P), or likely (LP) pathogenic (**Figure S3A**). These variants were very rare in the population as the total number of counts in the population was 457 including both homoplasmic and heteroplasmic calls (0.3% of the total counts of tRNA variants) (**Table S2**). Moreover, 56% of the calls for variants predicted to be of high-severity were heteroplasmic calls, which is significantly more compared to 27% for variants predicted to be of medium severity (p=1.3E-25, fisher-exact test), 2% for variants predicted to be of low severity (p=9.3E-280, fisher-exact test), and 22% for variants of unknown severity (p=4.4E-24, fisher-exact test) (**Figure S3B, Table S2**). There were 1,684 unique variants that mapped to the 2 rRNA genes (**Figure S3C-D**). The only annotation we were able to find to predict the impact of variants in rRNA genes was the heterologous inferential analysis (HIA) technique (Elson et al., 2015; Smith et al., 2014) (**Methods**). The application of this method is not yet fully automated, and we were only able to annotate a small number (∼3%) of the variants in rRNA genes (**Table S2**).

### Constrained regions in the mitochondrial genome

Inspired by the work to map the coding constrained regions in the nuclear genome (Havrilla et al., 2019), we looked for regions of the mitochondrial genome without any variation, hypothesizing that highly constrained regions may be functionally important. This may be particularly important for rRNA genes where very few annotations are available as evidenced in the previous section. When restricting our variant list to only homoplasmic calls, we observed that 7,938 bases were without any variation in this cohort (**Figure 5A**). When restricting to only homoplasmic calls plus heteroplasmic calls with an alternate read fraction (ARF) ≥0.5, we observed that 7,723 bases were invariable in this cohort (**Figure 5B, Table S3**). When considering all homoplasmic and heteroplasmic calls, we observed that 6,228 bases were invariable in this cohort (**Figure 5C**). The full lists of invariant bases for the model based on variants observed at homoplasmic levels or heteroplasmic levels with at least one individual with an ARF ≥0.5 + are reported in **Table S3**.

**Figure 5.**
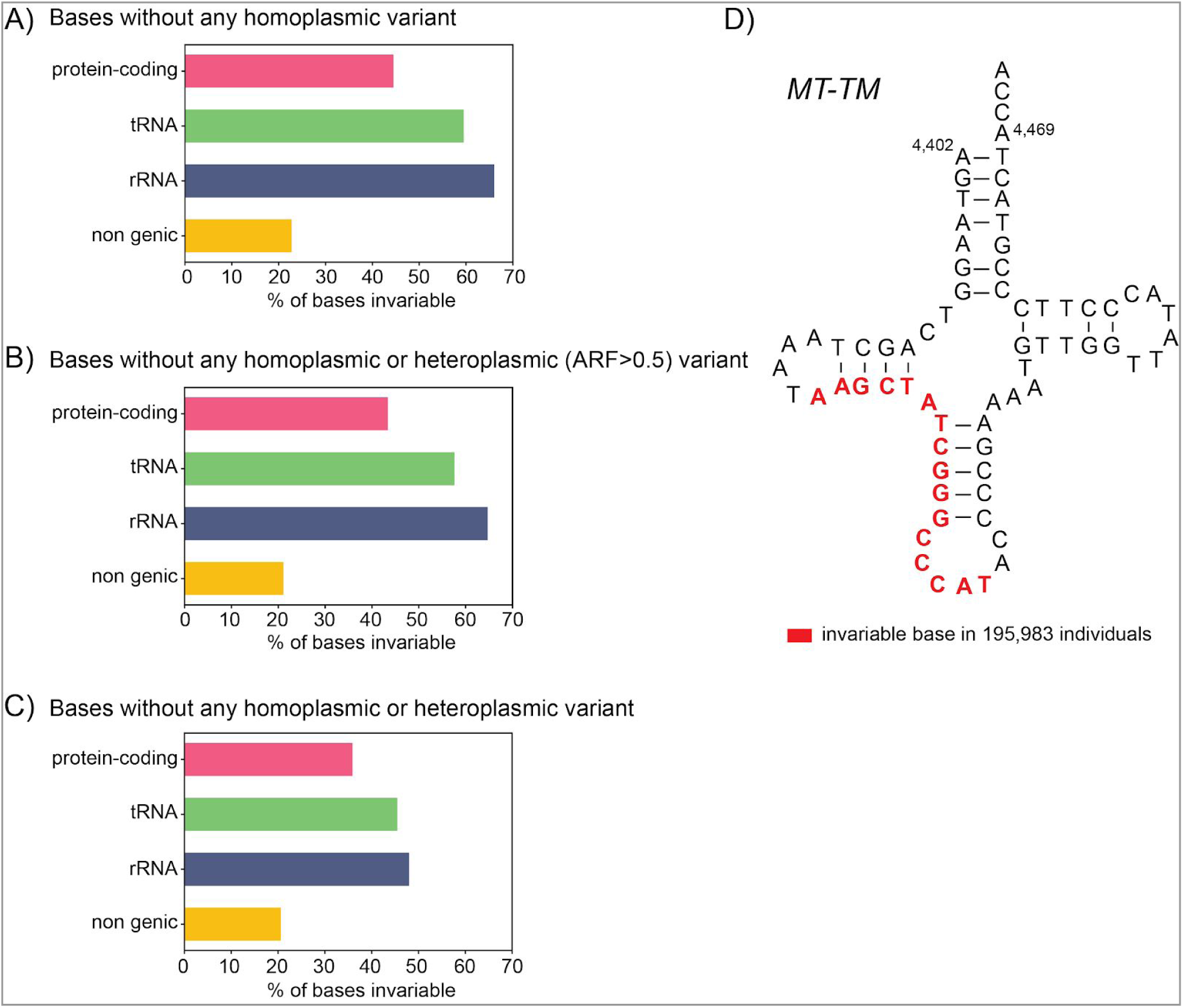
Constrained regions in the mitochondrial genome. **(A)** Proportion of the bases that were invariant when looking at homoplasmic variants only, grouped by genomic feature. **(B)** Proportion of the bases that were invariant when looking at homoplasmic calls plus heteroplasmic calls with an alternate read fraction (ARF) ≥0.5, grouped by genomic feature. **(C)** Proportion of the bases that were invariant when looking at all homoplasmic and heteroplasmic calls, grouped by genomic feature. **(D)** Visualization of a highly constrained region in *MT-TM*. Bases in red are bases that were invariable in the 195,983 mitochondrial genomes analyzed.

We then focused on the most constrained regions, which we defined as the longest stretches of mtDNA without any variation, when taking into account homoplasmic calls and heteroplasmic calls with a ARF≥0.5. We found 42 intervals of 11 bases or longer (**Table S3)**. We hypothesized that haplogroup markers should not be located within these constrained regions, and could be used as a control to verify the observed constraint. We obtained a list of 1,495 unique haplogroup markers from MITOMAP, using markers found at >=80% in haplogroups (Letter-Number-Letter). Indeed, we found that no haplogroup markers -- even those from haplogroups not represented in our dataset -- were mapped to these highly constrained regions (**Table S3**). In addition, no variants from PhyloTree Build 17 mapped to one of these highly constrained regions. Of note, the majority (28 out of 42) of these highly constrained regions were located in the 2 rRNA genes, and all but three of the remaining (11) were located in tRNAs (**Figure 5D**). This map of highly constrained regions will be helpful to decipher the role of specific domains of rRNA or tRNA genes, and will provide an additional annotation to interpret variants in noncoding regions, in tRNA and rRNA genes.

### Assessing classification of LHON variants reported in MITOMAP and ClinVar

In addition to identifying highly constrained regions that can help prioritize variants involved in severe developmental disorders, large databases with population allele frequencies of variants can help discriminate variants for researchers or physicians interested in rare diseases (even those with adult age of onset, or non-lethal phenotype). Inclusion in or exclusion from HelixMTdb was not based on any clinical phenotype. This database can therefore be used to assess whether a mtDNA variant is a good candidate variant for a rare mitochondrial disorder. If the frequency of a variant in HelixMTdb is above the maximum credible population allele frequency, then the variant is unlikely to cause a mitochondrial disorder by itself (**Figure 6A**). For a mitochondrial disorder assumed to be caused by a homoplasmic variant, the maximum credible population allele frequency can be calculated with the equation: *Maximum credible population AF = prevalence x maximum allelic contribution x 1/penetrance (Whiffin et al., 2017).* It is then possible to estimate the upper bound of the allele count expected in a population database given this maximum credible population AF (**Methods**).

**Figure 6.**
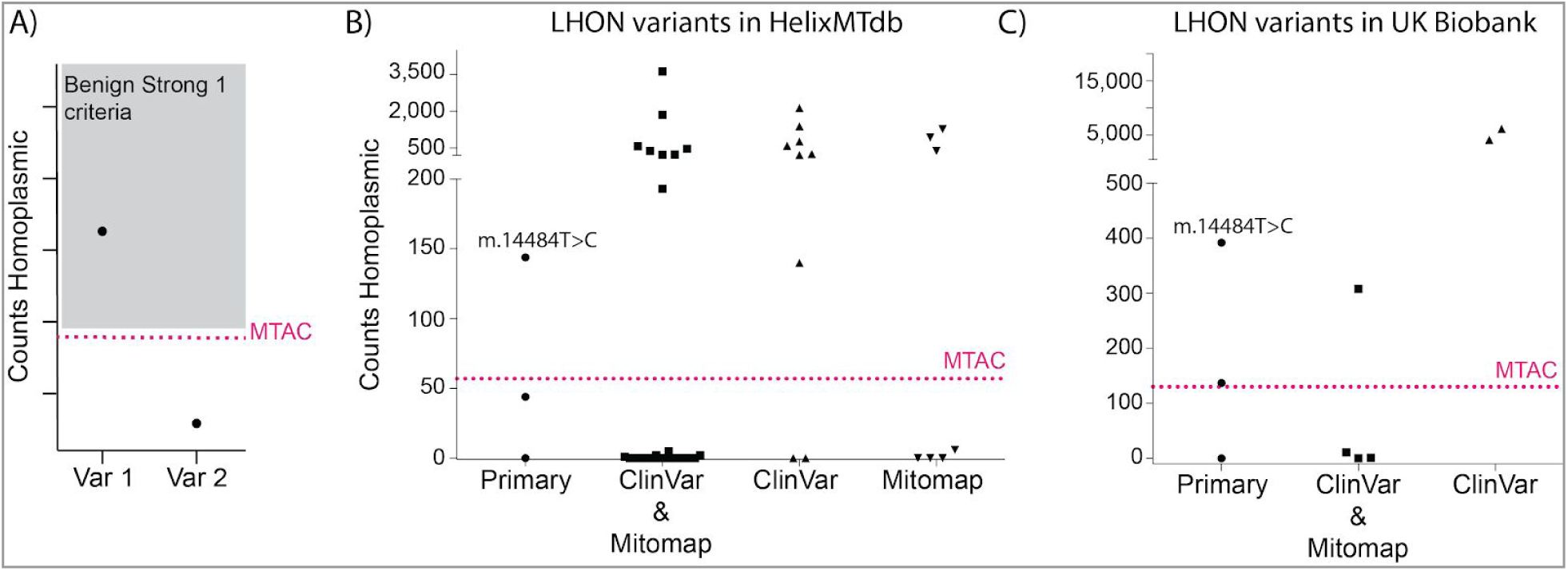
Counts of LHON variants in HelixMTdb and UK Biobank. **(A)** Visual aid to read the graphs and to assess whether the population allele frequency (AF) of a variant is higher than expected for a variant causing the disease, given what is known about the prevalence and the genetic architecture of the disease. MTAC is the Maximum Tolerated Allele Count (given a disease, the prevalence and genetic architecture of the disease, and the size of the database). The grey box indicates the zone where variants would meet the BS1 (Benign Strong 1) criteria defined by the American College of Medical Genetics and Genomics. The BS1 criteria provides strong evidence that variants in this zone would not be pathogenic / having a high impact on the disease studied. **(B)** Allele counts for reported LHON variants in HelixMTdb. Each circle, square or triangle represents a unique mitochondrial DNA variant. The 3 LHON primary mutations are represented by circles. LHON variants reported as pathogenic in ClinVar and present in the LHON MITOMAP page are represented by squares. Triangles represent LHON variants described as pathogenic on ClinVar or in the MITOMAP LHON page, but not by both. The pink dotted line represents the maximum tolerated allele count (MTAC), which is 56 for HelixMTdb. **(C)** Allele counts for reported LHON variants in UK Biobank. MTAC is 130 for the UK Biobank.

We tested the utility of HelixMTdb using this approach on Leber’s Hereditary Optic Neuropathy (LHON), which is one of the most studied mitochondrial disorders, with many references available to calculate the prevalence of the disease, genetic homogeneity, and penetrance (Yu-Wai-Man and Chinnery, 2000). With a model aimed at providing an upper estimate of the allele frequency in the population, we estimated the *LHON maximum credible population AF = 1/30,000 × 0.7 × 1/(0.1) = 0.00023*. Assuming that the number of observed variant instances in HelixMTdb follows a Poisson distribution (Whiffin et al., 2017), the expected allele count for LHON in HelixMTdb is 45, based on 195,983 mitochondrial genomes, with a maximum tolerated allele count (MTAC) for an LHON-causing variant of 56 (95% confidence). Variants reported to be LHON-causing in the literature should have allele counts below the maximum tolerated allele count calculated.

MITOMAP and ClinVar are two databases that catalog variants reported to be pathogenic for many mitochondrial diseases. As of July 2019, there were a total of 45 variants linked to LHON in either MITOMAP or ClinVar (**Table S4**). We grouped these variants based on the amount of evidence that supported the impact of the variant for LHON: (i) the 3 primary LHON variants, (ii) 26 additional variants reported as pathogenic in ClinVar and linked to LHON in MITOMAP, (iii) 9 variants reported as pathogenic in ClinVar but not reported in MITOMAP, and (iv) 7 variants on the MITOMAP LHON page (www.mitomap.org/foswiki/bin/view/MITOMAP/MutationsLHON, accessed in May 2020), but not reported as pathogenic in ClinVar (**Table S4**). We compared the observed counts for homoplasmic calls for these known LHON variants in HelixMTdb to the MTAC (summarized with their quality metrics in **Table S4)**. On average, the read depth (DP) was 168, the genotype quality (GQ) was above 95, the mapping quality (MQ) was 60 (see **Methods**), and the strand odds ratio (SOR) was 0.83, which altogether indicate that the calls for LHON variants were of high quality. Homoplasmic counts in HelixMTdb were above the MTAC for LHON for 19 (of 45) reportedly pathogenic LHON variants (**Figure 6B, Table S4**). These 19 variants are unlikely to be pathogenic by themselves, assuming the estimates regarding the prevalence of LHON, genetic homogeneity, and penetrance of the most common LHON variant are accurate.

One example of a variant whose frequency in this unselected cohort challenges existing literature is m.14484T>C, one of the three primary mutations for LHON (Brown et al., 1995, 1997; Torroni et al., 1997). This variant was present in 170 individuals, with 144 homoplasmic calls and 26 heteroplasmic calls, out of 195,983 individuals (AF_hom_: ∼9 in 10,000). Electronic medical records (EMR) were available for 18 of the 144 individuals with a homoplasmic m.14484T>C call in HelixMTdb. None of these 18 individuals had an ICD10 code starting with H47.2 in their electronic health record, which represents all optic atrophies, including hereditary optic atrophy (code H47.22) (**Table 1**). We then tested whether this result would replicate by looking at the allele frequency of m.14484T>C in the UK Biobank (UKB) cohort. The m.14484T>C variant and 8 other known LHON variants were directly genotyped with the UKB genotyping array. The allele frequency was AF_hom_: ∼8 in 10,000 in the entire cohort (n=392 individuals out of 486,036), and it was AF_hom_: ∼9 in 10,000 in a subset of unrelated individuals of European ancestry (n=291 individuals out of 335,840). These results confirmed the relatively high frequency of m.14484T>C variant in the population (**Figure 6C, Table S4**). Looking at the ICD10 codes in all UKB medical records, 97 participants had at least one ICD10 code H47.2 in their health records (optic atrophies have been recorded); however, none of the participants with the m.14484T>C variant had an ICD10 code starting with H47.2 (**Table 1**). Altogether, these analyses strongly suggest that the m.14484T>C variant does not cause LHON by itself.

**Table 1:** Phenotype of individuals carrying the m.14484T>C variant.

|  | HelixMTdb<br>all individuals | HelixMTdb<br>homoplasmic<br>m.14484T>C | UK Biobank<br>all individuals | UK Biobank<br>homoplasmic<br>m.14484T>C |
| --- | --- | --- | --- | --- |
| n samples | 195,983 | 144 | 486,428 | 392 |
| n samples with EHR available (% of samples) | 18,503 (9%) | 18 | 413,647 | 318 |
| Mean number of records in EHR when available (range) | 54 (1 - 823) | 39 (1 - 156) | 21 (1 - 5012) | 17 (1 - 209) |
| Median number of records in EHR when available | 35 | 34 | 8 | 8 |
| Number of individuals with ICD10 code H47.2 (% of samples) | 12 (0.06%) | 0 (0%) | 97 (0.02%) | 0 (0%) |

## Discussion

Here we present a genomic resource that can be used to answer new biological questions, and decipher the genetic etiology of rare mitochondrial disorders. HelixMTdb reflects the aggregated and de-identified mitochondrial DNA variants of 195,983 unrelated individuals. This is approximately 4 times more full mitochondrial genomes than what is currently available in MITOMAP or HmtDB (Lott et al., 2013; Preste et al., 2019), two prominent mtDNA variant databases. Unique properties of HelixMTdb are that: (i) it is not enriched for patients with mitochondrial disorders; (ii) it is less prone to batch effects since all samples were processed through the same lab protocol and variant calling pipeline; and (iii) it includes heteroplasmic calls and statistics on the allele fraction for these calls. It is also worth noting that all individuals sequenced were adults, with a median age group of 46-50, which is essential when evaluating candidate variants for rare and life-threatening diseases in childhood. This resource thus addresses the three main limitations of current population allele frequency databases for mtDNA variants, but also has its own limitations. A first limitation is that the average read depth per sample was 182 for the mitochondrial genome, which reduced the sensitivity for extremely low-fraction heteroplasmies (<10%). This limitation should not have an impact for the use of the database in a clinical setting as the majority of pathogenic variants have a functional impact when heteroplasmy levels are >70% (Russell et al., 2020). A second limitation comes from the fact that mitochondrial DNA in this study was extracted from saliva, and heteroplasmic levels may not reflect levels present in mitochondria from phenotype-affected tissues such as muscle. A third consideration is the relatively low diversity of mitochondrial genomes / haplogroups represented in HelixMTdb. For example, a smaller percentage of the individuals came from the L lineages (African) or M lineages (Asian) in HelixMTdb compared to MITOMAP.

We identified 14,324 unique variants, excluding variants overlapping homopolymer tracts. We showed that 20% of the variants were only observed at heteroplasmic levels, which would be missed if heteroplasmic calls were not included. When looking at the 13 protein-coding genes, we showed that the mitochondrial genome is not tolerant to protein-truncating variants at high levels of heteroplasmy. This is in contrast with the relative tolerance of the mitochondrial genome for missense variants as shown by the high number of missense variants observed at homoplasmic levels in the population, which was very close to the number of synonymous variants observed at homoplasmic levels in the population. The only exception to this rule was the presence of a nonsense variant at the start codon of *MT-ND1*. All of the individuals carrying the nonsense variant were from the same haplogroup T. It is also present in MITOMAP in individuals from the T haplogroup. It is likely that MT-ND1 is still properly translated in these individuals with the use of a non-canonical start codon in the context of the T haplogroup, or with the use of the Methionine encoded at the third cdon.

We found that 47% of the bases of the mitochondrial genome did not even have one homoplasmic or heteroplasmic call at a level higher than 50% across the entire cohort. Given the fact that the number of unrelated individuals in the cohort was >10x the number of bases in the mitochondrial genome, and the fact that the mutation rate of mtDNA is higher than the mutation rate of nuclear DNA (Sigurðardóttir et al., 2000), this result shows the very high constraint on the mitochondrial genome. Notably, the two rRNA genes were under highest constraint with 65% of their bases invariant. This high level of constraint is potentially the result of the absence of redundancy for the mitochondrial rRNA genes, unlike the rRNA genes in the nuclear genome that are present in >100 copies located in five rDNA clusters (Lander et al., 2001). Most of the known modifications of 16S rRNA and 12S rRNA fall within the most highly constrained regions (Hällberg and Larsson, 2014). The tRNA genes also showed strong constraints, and the smaller representation of the tRNAs in the longest stretches without any variant may be explained by the smaller size of tRNA genes compared to the 2 rRNA genes. At the opposite of rRNA and tRNA genes, most of the non-coding bases were variable in the population. The one exception to this rule was a short stretch mapping to the mitochondrial light strand origin of replication.

We hope that these maps of highly constrained regions in the mitochondrial genome will be used to annotate variants, especially those falling in rRNA genes. Here are three illustrations. First example: the regions under high constraint in the rRNA genes allow molecular biologists to design experiments to study translation and regulation of protein expression in the mitochondria (Hällberg and Larsson, 2014). Second example: these annotations could be used to analyze somatic mitochondrial mutations identified in cancers (Reznik et al., 2017; Yuan et al., 2020). For example, a recent study reported a ‘comprehensive’ molecular characterization of mitochondrial genomes in human cancers based on 2,658 cancers from The Cancer Genome Atlas, but the study focused on the impact of truncating variants (and other nonsynonymous variants) as well as mitochondria copy number (Yuan et al., 2020). The analysis of variants in rRNA genes was omitted possibly due to the difficulty of analyzing these without available annotations. We hope HeixMTdb will help in that context. Third example: this is useful when interpreting the potential role of a variant for a rare (and potentially life-threatening in childhood) developmental disorder. Variants in these regions under high constraint are very good candidates.

We have also shown that this resource can be used to better evaluate and prioritize variants suspected to cause rare mitochondrial disease, as long as some assumptions on the prevalence, and genetic architecture of the disease could be made. Through a comparison of the allele frequencies of LHON variants to disease prevalence, we showed that ∼40% of the variants reported to be pathogenic for LHON on ClinVar or MITOMAP could be re-classified as Benign / Likely Benign based on the ACMG standards and guidelines because the Benign Strong 1 (BS1) criteria would apply (Richards et al., 2015; Wong et al., 2020). In particular, the primary LHON m.14484T>C variant is likely not pathogenic for LHON by itself. We were able to replicate these results using the UK Biobank cohort, and we showed that the frequency of the variant in unselected cohorts is high, with very low penetrance (0/144 and 0/392 individuals had a LHON diagnosis in their health record). These results are consistent with previously reported pedigree analyses finding that this variant exhibits a low LHON penetrance in a non-haplogroup-J background (Brown et al., 1997; Howell et al., 2003; Puomila et al., 2007; Torroni et al., 1997). Of note, the m.14484T>C variant was present in 13 different haplogroup lineages in HelixMTdb (**Table S4**), and the ratio of (haplogroup J m.14484T>C carriers) / (all haplogroup J) = 4 / 16,030 was the lowest compared to the ratio for the 12 other haplogroups. It remains a possibility that there may be a branch of haplogroup J where a combination of variants with m.14484T>C is pathogenic (Brown et al., 1995; Carelli et al., 2006). Overall, our analysis of variants reported to be pathogenic in ClinVar and MITOMAP for a well-characterized mitochondrial disorder highlights the clinical utility of HelixMTdb. We believe this resource will be instrumental in improving clinical classification of variants, similar to the role that other large nuclear DNA variation databases play in clinical interpretation today (Richards et al., 2015).

## Acknowledgements

We thank all Helix users, all participants in the Healthy Nevada Project, as well as all research participants in the UK Biobank project. This research has been conducted using the UK Biobank Resource under Application Number 40436. We acknowledge Dr. Ekaterina Yonova-Doing for initial work on the frequency of the m.14484T>C variant in the UK Biobank. We also thank Dr. Agnel Sfeir and Dr. Patrick Chinnery for discussions about mitochondrial biology and about ideas of experiments to perform. We acknowledge the Helix Laboratory Operations team as well as the Bioinformatics team for their contributions to the production of clinical-grade exomes, the downstream QC and analysis pipelines. We thank the early users of this resource who have provided great feedback and ideas on how to improve the resource, especially the Mitomap team.

## Declaration of Interests

AB, FM, SW, FJ, MI, RJ, ADR, EC, MR, WL, JL and NW are employees of Helix.

## Methods

### Individuals

The HelixMTdb database reflects aggregated and de-identified mitochondrial DNA variants observed in individuals sequenced at Helix. The cohort is skewed slightly female at 52%, with a non-normal distribution of samples aged 18-85+ (median age group = 46-50). All individuals sequenced resided in the United States at the time of providing their saliva sample. Importantly, these individuals have not been sequenced based on the presence or absence of any medical phenotype (i.e. there are no inclusion or exclusion criteria in the registration process based on any medical phenotype). Nine percent of Helix users in this study were also participants in the Healthy Nevada Project under the University of Nevada Reno IRB protocol: #7701703417. Electronic medical records were available for most of the Healthy Nevada Project participants, and these records showed no enrichment for classic mitochondrial diseases as shown in **Table S5**.

The replication study for the primary LHON variants was based on the UK Biobank resource (Sudlow et al., 2015), under application number 40436.

### Sample preparation, Sequencing, and Variant Calling

Library Preparation and Enrichment was performed in the Helix clinical laboratory (CLIA #05D2117342, CAP #9382893). Samples were sequenced using the Exome+ assay, a proprietary exome that combines a highly performant medical exome, the mitochondrial genome, and a microarray-equivalent SNP backbone into a single sequencing assay (www.helix.com). Read length was 75 bp. Base calling and alignment were run on BaseSpace servers. For mitochondria, we first extracted read pairs in which both reads were mapped and at least one was mapped to the mtDNA. This enabled us to map regions that might otherwise be discarded due to multimapping regions of homology with nuclear sites (NUMTs). Reads were mapped to the rCRS (GenBank: J01415.2) using BWA mem (Li, 2013), and were deduplicated and realigned using the Sentieon implementation of the GATK algorithms(DePristo et al., 2011) (Freed et al., 2017). VCF files were generated using haplotyper with `emit_mode=confident`.

The mean read depth across the mitochondria for an individual was DP=182.

Before including samples and calls into HelixMTdb, we used the following filters: removed samples with mean mtDNA coverage <20; removed calls at positions covered by less than 10 reads; removed calls with genotype quality (GQ) below 20.

### Haplogroup Calling

We collapsed heteroplasmic calls into either ALT or REF homoplasmic calls whenever the majority call consisted of at least 75% of the total reads. The remaining sites were left as heteroplasmic, although they are ignored (assumed as reference) by the haplogroup caller. Some indels in a VCF can be left- or right-aligned, meaning that they could be expressed in more than one fashion, changing the coordinates of the positions that are affected. For instance, a change in the number of repeats of a microsatellite can be expressed as an indel at the beginning or at the end of the microsatellite. Both of these can be used to reconstruct the same sequence, but might be used differently by haplogroup callers. Our pipeline originally makes a left-alignment, which is the way the calls are represented in HelixMTdb. We changed it to a right-alignment to be able to use Haplogrep (Weissensteiner et al., 2016). We removed, in advance, sites and mutations that were not incorporated in Phylotree v17 (van Oven and Kayser, 2009), because they are not commonly used for haplogroup assignment. We then ran Haplogrep, using rCRS as reference, and kept the first (ranked) 40 hits for further analysis. A number of steps were taken to further reduce the number of haplogroups under consideration: (i) the quality call (for haplogroup) had to be at least 0.94 of that of the maximum, and (ii) at least as high as the value of the third ranked quality (ties might result in more than three haplogroup passing this filter). The most recent common ancestor of these haplogroups was selected as representative for the sample. When the lineage falls in a haplogroup that is similar to that of rCRS, Haplogrep tends to provide inaccurate results. For instance, a VCF file without any position, does not provide a haplogrep call, but corresponds to a sequence that matches the rCRS. Also, lineages similar to the rCRS that share a mutation with a different part of the tree might be assigned to an incorrect haplogroup, but are characterized by a large number of missing mutations for the haplogroup. These were all corrected afterwards.

For comparative representation in HelixMTdb, we combined haplogroups into higher-level haplogroups that matched those shown in MITOMAP (**Table S1**). For HelixMTdb, we further grouped higher-level haplogroups with less than 10 individuals with other higher-level haplogroups to avoid providing an individual’s full mitochondrial DNA sequence. This resulted in the grouping together of ‘L4 + L5 + L6’ and ‘X + S’.

### Relatedness analysis

In addition to calling mitochondrial DNA variants, reads from the entire Exome+ were mapped to Human Reference GRCh38 for non-mitochondrial variant calling, using a custom version of the Sentieon align and calling algorithms (Kendig et al., 2018) following GATK best practices. For allele frequency analysis, we further reduced the sample set by removing individuals related at the 2nd-degree or closer.

Briefly, we calculated kinship using the Hail `pc_relate` method ([CSL STYLE ERROR: reference with no printed form.]) using 11,772 representative common SNPs spread across the genome. The method `pc_relate` was run with the first 10 principal components and a kinship cutoff of 0.0884.

From clusters of family members, we kept both halves of the father-child relationships. For other relationships, we randomly selected one representative to retain. In total, 21,074 samples were removed at this stage. We labeled this the unrelated dataset, and proceeded with our analysis based on this cohort of 195,983 individuals.

### Ancestry assignment and principal component analysis

For each individual we ran a supervised ADMIXTURE algorithm with k=5 ancestral populations. From these admixture coefficients, we then labeled each individual with one ancestry using the following decision tree:

- When (ADMIX_EUR>0.85) & (ADMIX_EAS<0.1) & (ADMIX_SAS<0.1) & (ADMIX_AFR<0.1) & (ADMIX_AMR<0.1) then “European”
- When ADMIX_EAS>0.6 then “East Asian”
- When ADMIX_SAS>0.6 then “South Asian”
- When (ADMIX_AFR>0.3) & (ADMIX_EAS<0.1) & (ADMIX_SAS<0.1) & (ADMIX_AFR > ADMIX_AMR) then “African”
- When (ADMIX_AMR>0.1) & (ADMIX_EAS<0.1) & (ADMIX_SAS<0.1) then “Latinx)
- Default (“Other”)

Principal Component Analysis was done using Hail `hwe_normalized_pca` function and based on 12,000 autosomal SNPs spread across the genome.

### Analysis of Allele Frequency and Heteroplasmy

All allele frequency and heteroplasmy analysis, as well as all of the analyses listed after this paragraph, were performed in Hail (<u>Hail Team. Hail 0.2.13-81ab564db2b4</u>. https://github.com/hail-is/hail/releases/tag/0.2.13.) on Amazon HPC clusters. Briefly, batches of 500 gVCF files were combined into multi-sample gVCF files using GenomicsDB (https://github.com/Intel-HLS/GenomicsDB/wiki). Multi-sample VCF files were extracted from the resulting gVCF at sites deemed to be informative and then combined into large pVCF files for ingest into Hail using Bcftools (https://samtools.github.io/bcftools/bcftools.html). All variants were left-aligned.

Levels of heteroplasmy play important roles in causing a mitochondrial disease, as well as modulating the strength of phenotypes. To provide as much information as possible regarding the levels of heteroplasmy observed for each heteroplasmic call, we defined *ARF = Alternate Read Fraction = (counts of reads supporting the alternate allele) / (count of all reads at this position).*

### Comparison with MITOMAP database

We downloaded the MITOMAP GenBank FL ID set (all of the full-length sequences) on June 16, 2019 (http://www.mitomap.org). The MITOMAP database at the time was based on 47,412 full-length mitochondrial sequences (www.mitomap.org/foswiki/bin/view/MITOMAP/Mitobank). However, the maximum allele count displayed for one variant was 48,241. For the Mitomap Allele Frequency calculation, we considered that there were 48,241 mitochondrial genomes with coverage at every base.

We compared the variants, their counts and their allele frequencies in multiple ways. We looked at all calls, homoplasmic SNVs, homoplasmic insertions, and homoplasmic deletions. We plotted the results using a scatter plot, and calculated the Spearman rho coefficient. The most notable differences in variant calls between HelixMTdb and MITOMAP were observed in the homopolymer stretch between position m.302 and m.315. We think that it is likely that both HelixMTdb and MITOMAP have inaccurate calls at this locus. In addition, some differences in variant frequencies between the two databases may be due to differences in left- or right-alignment in homopolymer stretches.

### Annotation of feature type

Genomic feature locations were annotated using the list from MITOMAP (https://www.mitomap.org/foswiki/bin/view/MITOMAP/GenomeLoci), and further curated into four groups: protein-coding, rRNA, tRNA, and non-coding (all remaining sites including the D-loop).

Moreover, a few positions overlap multiple features (e.g. positions 4329-4331 overlapping *MT-TI* and *MT-TQ*, or positions 5721-5729 overlapping MT-TN and the noncoding L strand origin MT-OLR). In these cases, we made arbitrary decisions to avoid overlapping annotations that may impact some future analyses. The positions and their associated feature type are represented in **Table S6**.

### List of constrained intervals in the mitochondrial genome

To calculate invariable positions in HelixMTdb, we defined a position as being variable if at least one SNV, or one deletion was overlapping this position. **Figure 5** provides the results taking into account (A) only homoplasmic calls, or (B) homoplasmic calls and heteroplasmic variants where at least one individual was observed with a ARF ≥0.5, or (C) all homoplasmic and heteroplasmic calls. We used BEDTools (Quinlan and Hall, 2010) to `sort` and `merge` the list of SNVs and positions deleted, and defined the final list of positions that were variable.

We then used `bedtools complement` to obtain the list of constrained intervals in the mitochondrial genome.

### PhyloTree variants in highly constrained intervals

To test that the regions identified as highly constrained are invariable in the main structure of the mitochondrial phylogenetic tree, we collected the list of mutations from the official <u>phylotree</u> page. After trimming the characters that do not identify position (e.g. ref base, their character of recurrent, deletion, insertion), we generated a list of all positions, and a BED file spanning those positions. Likewise, we generated a BED file for the intervals indicated in **Table S3**. Using BEDTools we assessed the intersection of these BED files, and the result is that the intersection was empty.

### Annotation of impact and predicted severity

All variants were classified using Variant Effect Predictor (VEP) against ENSEMBL e!95. Conservation scores were reported from phastCons 100way_vertebrate, obtained from UCSC for GRCh38 (http://hgdownload.cse.ucsc.edu/goldenpath/hg38/phastCons100way/hg38.100way.phastCons/chrM.phastCons100way.wigFix.gz).

For protein-coding variants, severity was determined using the VEP `most_severe_consequence` annotation, and grouped as follows in **Table S2**:

- High: frameshift, stop_gained, stop_loss
- Medium: inframe indel, missense, start loss
- Low: synonymous, stop_retained

tRNA variants were classified by submitting variants in tabular format to the Mitomaster Web Service API, obtaining their raw MitoTip (Sonney et al., 2017) score, then converting the score to a predicted pathogenicity, using the MitoTip scoring matrix (https://www.mitomap.org/foswiki/bin/view/MITOMAP/MitoTipInfo) as follows: >16.25=Likely Pathogenic (LP); 12.66-16.25=Possibly Pathogenic (PP); 8.44-12.66=Possibly Benign (PB); <8.44=Likely Benign (LB). Mitotip tRNA pathogenicity scores are a result of a combination of conservation, frequency in databases, and predicted secondary structure disruption. In **Table S2**, these tRNA variants were combined as follows:

- High severity: known Pathogenic (P), likely pathogenic (LP)
- Medium severity: Possibly pathogenic (PP)
- Low severity: Benign (B), likely benign (LB), and possibly benign (PB)

rRNA variants were annotated using the list of variants and their severity categories determined by Heterologous Inferential Analysis (HIA) published in (Elson et al., 2015; Smith et al., 2014). Briefly, this technique maps rRNA variants onto the crystal structure for Human 12S and 16S subunits to understand likely structural defects and leverages functional assay results from highly conserved homologs in multiple species to assign pathogenicity. Of the 113 variants derived from Genbank sequences collected from these two papers, we were able to annotate the predicted severity for only 43 matching variants. In addition, we found 1,807 novel variants that have not been previously classified in the literature or reported in MITOMAP. We left these as “Unknown” severity.

### Maximum tolerated allele count

Our main objective was to filter out variants that could not be disease-causing given a pre-defined genetic architecture. The objective was not to prove the pathogenicity of any given variant. Therefore, we opted for a conservative model that would minimize the number of variants discarded and provide a high estimate of the maximum credible population AF. The method and calculations used here are almost identical to a method previously published to calculate the maximum credible population AF, and maximum tolerated allele count for dominant disorders (Whiffin et al., 2017)

#### Maximum credible population AF = prevalence x maximum allelic contribution x 1/penetrance

For LHON:

- <u>Genetic architecture</u>: disease is caused by a <u>homoplasmic</u> mtDNA variant.
- <u>Prevalence in the population: 1 in 30,000</u>. Reports have shown that prevalence is about 1/31,000 in the North East of England, and 1 in 50,000 in Finland (Puomil et al., 2007; Yu-Wai-Man and Chinnery, 2000; Yu-Wai-Man et al., 2003).
- <u>Maximum allelic contribution: 0.7</u>. The three primary LHON mutations (m.3460G>A, m.11778G>A, and m.14484T>C) explain the majority of reported LHON cases. Among these, the m.11778G>A is accounting for approximately 70% of cases among northern European populations (Mackey et al., 1996; Yu-Wai-Man and Chinnery, 2000). Overall, we felt like one variant accounting for 70% of LHON cases in our cohort from all parts of the United States was a very high estimate for maximum allelic contribution.
- <u>Penetrance: 0.1</u>. Penetrance is probably the harder number to estimate for this equation. Of note, the penetrance for LHON is sex-specific (Yu-Wai-Man and Chinnery, 2000). Males have a much higher risk of developing symptoms than females. The ranges of the risk of developing symptoms were 32-57% for males and 8-28% for females (Yu-Wai-Man and Chinnery, 2000; Yu-Wai-Man et al., 2003). To be conservative, we selected a penetrance number on the lower end of these ranges.
- <u>Result:</u> **Maximum credible population AF for LHON = 0.00023**.

To calculate the maximum tolerated allele count (MTAC), we calculated the allele count at the upper bound of the one-tailed 95% confidence interval for the established maximum allele frequency, given the number of alleles in the population database. An approximation using a Poisson distribution has been previously reported (Whiffin et al., 2017), and we used the same method in R.

#### MTAC = qpois(quantile_limit, an*af)

where *an* is the number of total alleles in the database, and *af* is the maximum credible population allele frequency.

For LHON in HelixMTdb:

~~~
MTAC = qpois(0.95, 195983*0.00023) = 56
~~~

We also looked at LHON variants in the UK Biobank cohort. Genotyping information was available for 265 mtDNA positions, for 488,377 samples.

For LHON in UK Biobank:

~~~
MTAC = qpois(0.95, 488377*0.00023) = 130
~~~

### LHON variants

The list of LHON variants from MITOMAP was copied and pasted in July 2019 from this address: https://www.mitomap.org/foswiki/bin/view/MITOMAP/MutationsLHON. The list of LHON variants from ClinVar was obtained using the following steps:

- Started from clinvar_20190603.vcf.gz (obtained here: ftp://ftp.ncbi.nlm.nih.gov/pub/clinvar/vcf_GRCh38/)
- Selected mitochondrial DNA variants
- Kept variants that included ‘Leber’s_optic_atrophy’ in the CLNDN field.
- Selected variants that were labeled as Pathogenic (of note, there were no Likely Pathogenic variants).

### Analysis of LHON phenotype in electronic medical records

Electronic medical records were analyzed by parsing the ICD10 codes. No filters were applied based on the source or date of entry. The code H47.2 was used for all Optic Atrophies, which includes the LHON phenotype H47.22). Other ICD10 codes related to eye diseases or other mitochondrial diseases were used as controls. Details are in **Table S5**.

## Data availability

This database is published under a <u>Creative Commons Attribution-NonCommercial-ShareAlike 4.0 License</u>, and may be used, shared and redistributed appropriately. Please cite this paper when using this database.

HelixMTdb can be downloaded using this link: https://s3.amazonaws.com/helix-research-public/mito/HelixMTdb_20200327.tsv

## Supplementary Information (3 supplementary figures and 6 supplementary tables)

**Figure S1 related to Figure 1.**
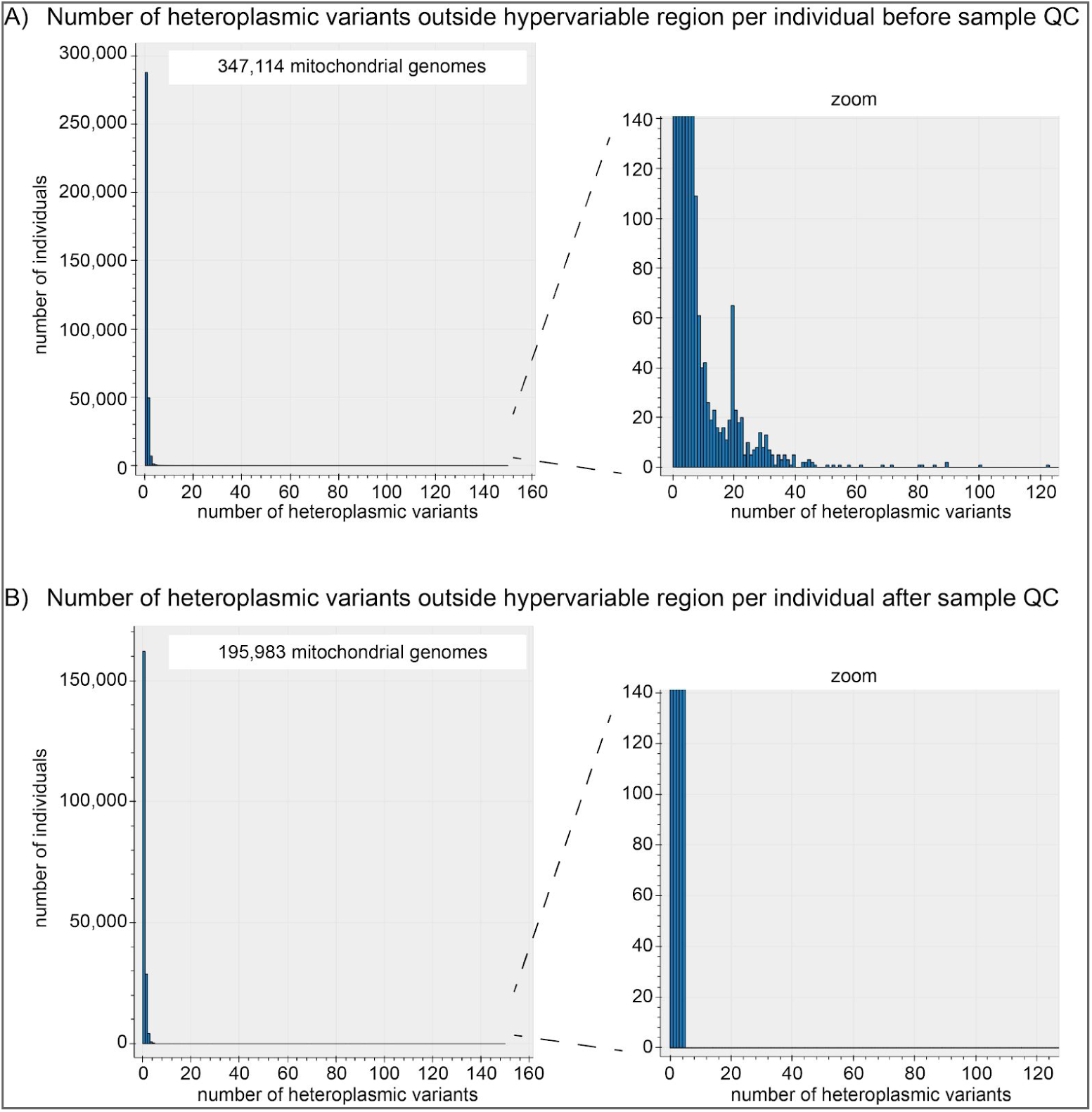
Overview of the 195,983 individuals and mitochondrial genomes aggregated in HelixMTdb. Distribution of the number of heteroplasmic SNVs outside of the hyper-variable region per individual. The panels on the right are a zoom of the panels on the left. **(A)** in the initial 347,114 mitochondrial genomes analyzed. **(B)** in the final 195,983 mitochondrial genomes included in HelixMTdb.

**Figure S2 related to Figure 2.**
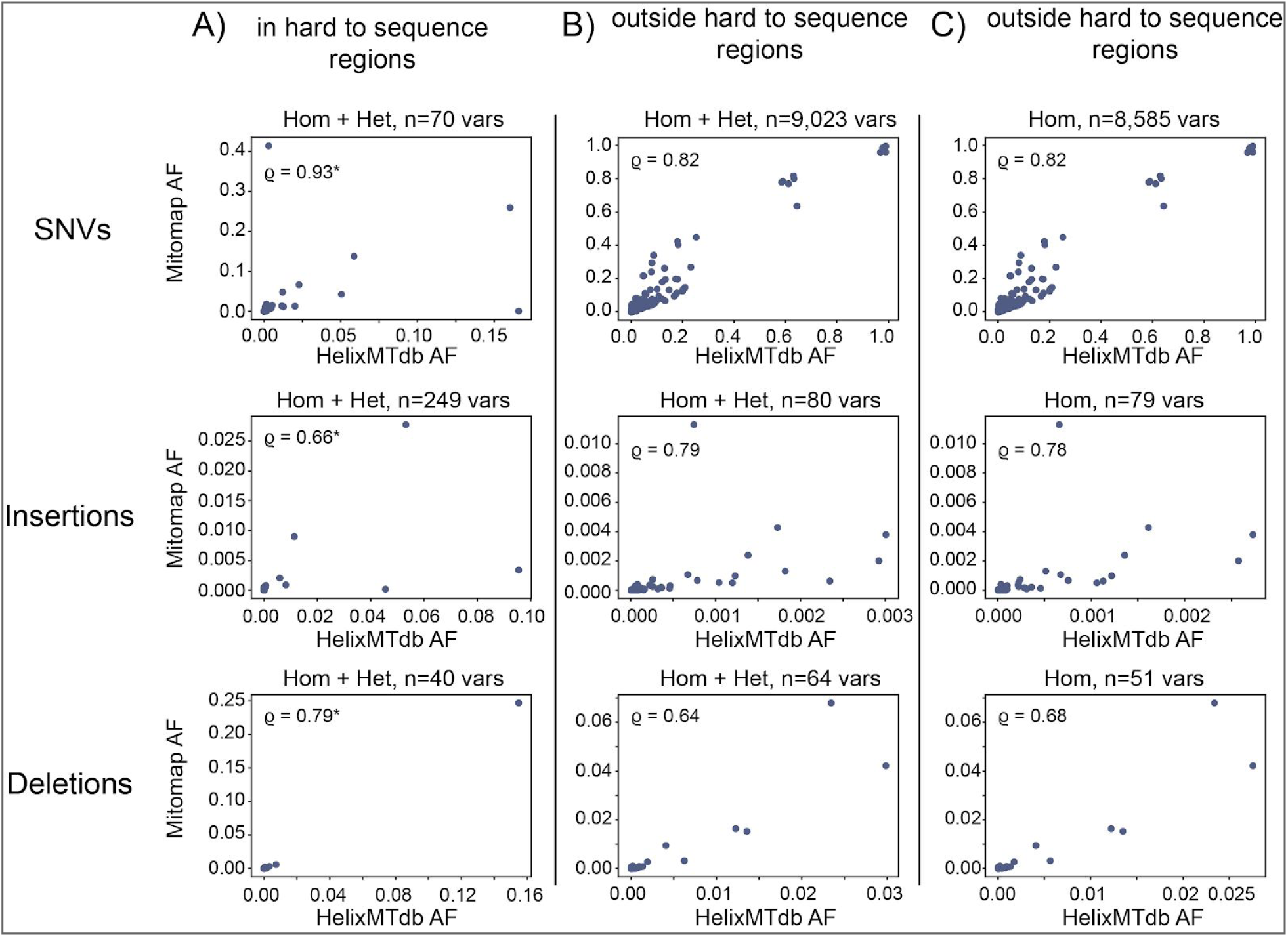
Comparison of the variants and their allele frequencies in HelixMTdb and MITOMAP. Graphs are scatter plots where each variant is represented by one dot. The x-axis represents AF in HelixMTdb, and the y-axis represented AF in MITOMAP. The spearman rho (ϱ) coefficient rho is indicated on the upper left of the plot. **(A)** AF of variants within the 3 hard-to-sequence regions: m.300-316, m.513-525, and m.16182-16194. **(B)** AF of homoplasmic and heteroplasmic variants in HelixMTdb, and all variants in MITOMAP -- excluding the hard-to-sequence regions -- are represented. **(C)** AF of homoplasmic variants in HelixMTdb, and all variants in MITOMAP -- excluding the hard-to-sequence regions -- are represented. **(A,B,C)** For each, the top graph represents SNVs, the middle graph represents insertions and the lower one represents deletions.

**Figure S3 related to Figure 4.**
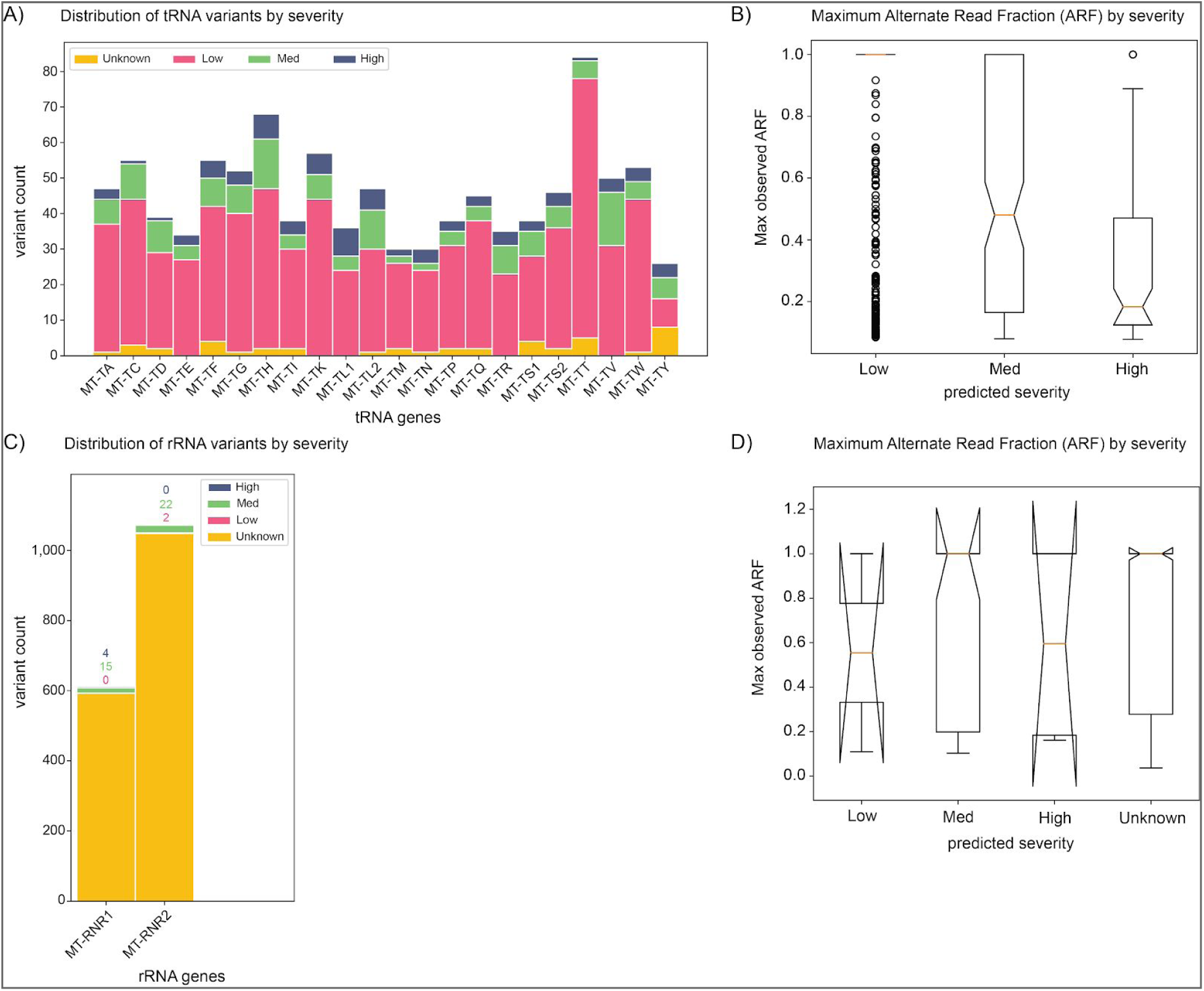
Intolerance to loss-of-function variants in tRNA and rRNA genes. **(A)** Summary counts of variants per tRNA gene, colored based on predicted severity. Pink: low; green: med; blue: high; yellow: unknown. Severity was calculated using MitoTip. **(B)** Distribution of the maximum observed Alternate Read Fraction (ARF) in tRNA genes for each severity category. **(C)** Summary counts of variants per rRNA gene, colored based on predicted severity. Pink: low; green: med; blue: high; yellow: unknown. Severity was manually determined from previous publications. **(D)** Distribution of the maximum observed Alternate Read Fraction (ARF) in rRNA genes for each severity category. Note the very low number of variants for the ‘Low’, ‘Med’ and ‘High’ groups creating these strange-looking Box plots.

**Table S1, related to Figure 1: Distribution of mitochondrial haplogroups in HelixMTdb**

**Table S2, related to Figure 4: Variant attributes by genomic features**

- variants in ‘hard to sequence’ regions were excluded.
- Mean allele frequency = ratio of n_non_ref / n_samples. So the % of individuals either with a homoplasmic or heteroplasmic variant vs number of total samples.
- % conservation = hl.agg.mean(phastcons100v).

**Table S3, related to Figure 5: List of all constrained mitochondrial regions inferred from homoplasmic calls and heteroplasmic calls with a ARF >=0.5**

Regions / intervals of 1bp were not included in this analysis.

**Table S4, related to Figure 6: Counts of all LHON variants reported in HelixMTdb and the UK Biobank**

**Table S5, related to Figure 6 and Table 1: ICD10 codes in Healthy Nevada Project and UK Biobank**

**Table S6, related to Methods: Location of genomic features in the mitochondrial genome**

## Notes

https://s3.amazonaws.com/helix-research-public/mito/HelixMTdb_20200327.tsv

### Summary of Updates

We added more stringent QC threshold for very low levels of contamination. These low levels of contamination were undetectable in the nuclear genome, but could have a small impact on heteroplasmic variants with low levels of heteroplasmy. This resulted in the removal of a couple hundreds samples and a few variants from the database. We provided an updated link to the database and have redone all of the analyses with the new samples and variants. We also added Figure 1 to better characterize the population. Figure 5 is also new.

